# Scalable prediction of symmetric protein complex structures

**DOI:** 10.1101/2025.11.14.688531

**Authors:** Victor Yu, Perry Demsko, Roger Castells-Graells, Hank Parker, Andrew Huang, Chloe Chen, Martin Huang, Varsitaa Srinivasan, Krishna Ajjarapu, Nica Tofighbakhsh, Richard Yu, Michael Lake, David L. Glanzman, Sarah Warren, Joseph Alzagatiti

## Abstract

All life relies on proteins to function, yet accurately modeling protein structures that exceed ≈ 10, 000 amino acids or have higher-order geometries remains difficult. Existing solutions are limited to specific scenarios, require considerable computational resources, or are otherwise unscalable. Consequently, many large, disease-relevant protein complexes in the human proteome, as well as nearly all viruses and numerous other classes, are impractical to model with high fidelity for drug development. To modulate these protein complexes and viruses, structural information is eminently valuable, and often essential. In the last two years, machine learning based-tools that can generate binders to a given target structure with high hit rates have emerged. Combined with high-throughput screening, these technologies can far outpace traditional drug discovery. However, they cannot function well without accurate models of their target structures. Thus, to unlock the full power of AI-driven drug discovery, a scalable method must be developed to predict large protein complex structures. To overcome this bottleneck, we introduce Plica-1, a physics-based method to rapidly and accurately predict the structure of arbitrarily large, symmetric protein complexes. Validated across 4 major symmetry classes (icosahedral, tetrahedral, octahedral, and cyclic), the method consistently achieves near-experimental levels of accuracy, i.e., RMSD < 5Å. In test cases, the method runs in < 5 minutes on consumer hardware, 10^3^-10^5^ times faster than the closest comparable software. The largest structure currently built, at ≈40,000 amino acids, is > 8 times the limit of existing machine learning methods. The results demonstrate that protein complexes can be modeled at significantly improved speeds and scales, making Plica-1 a promising tool for protein engineering and drug development.

## 1 Introduction and motivation

Developing a fast and accurate algorithm that can predict the structure of any protein assembly solely from its amino acid sequence is a central goal of structural biology.

Protein structures exist at three levels of scale: monomer, multimer, and complex (**Figure 1**). From the perspective of a protein complex, a single monomer corresponds to a brick; a single constituent multimer corresponds to a wall; and the entire complex, a house. Within *in vivo* systems, components of large protein complexes rarely assemble simultaneously, but rather in a sequential manner. Complexes often assemble monomers to multimers before fitting these multimers into the final assembly, prioritizing optimal interfaces between protein monomers before subsequent chains can be added [5, 6].

**Figure 1.**
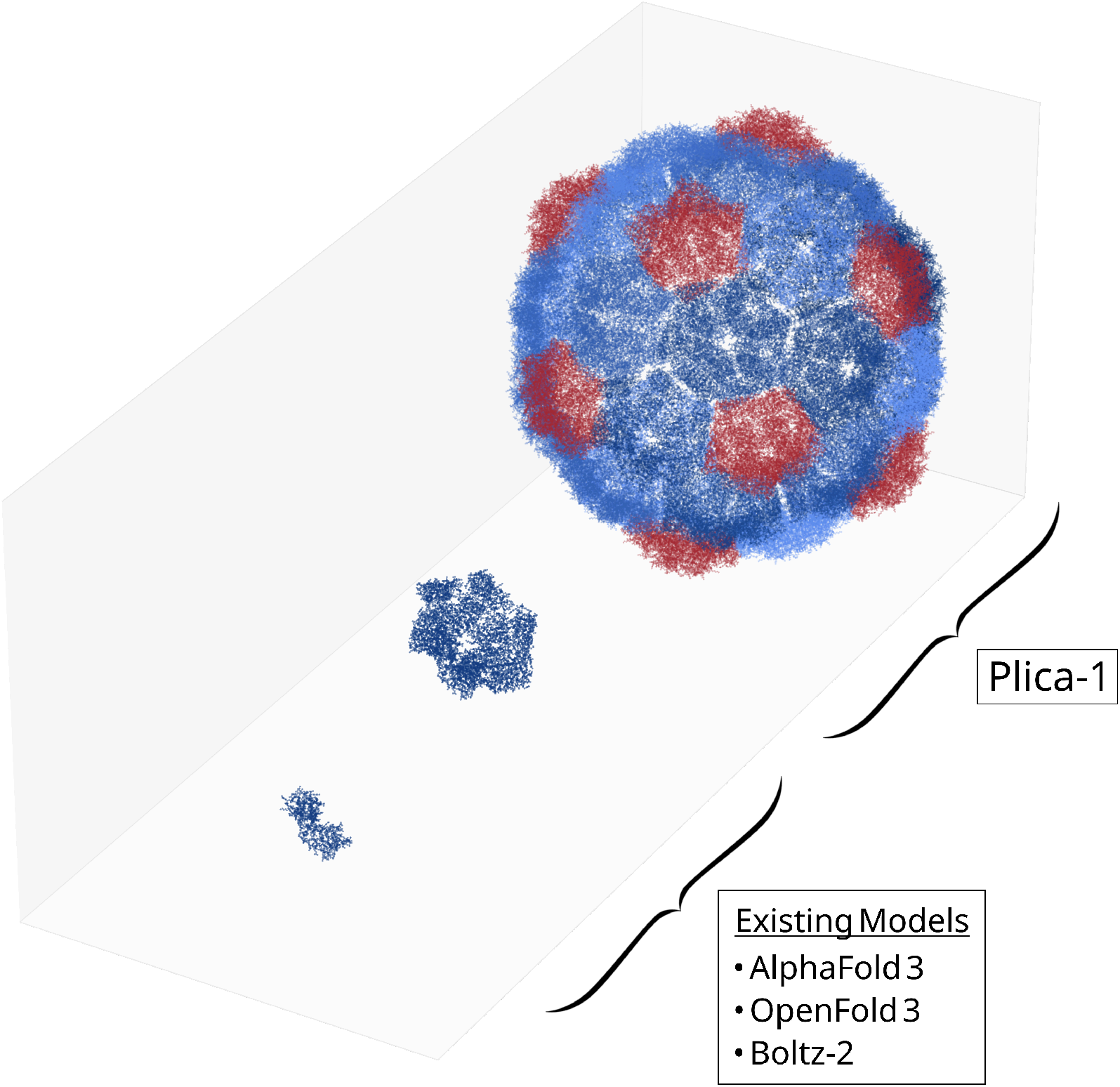
From left to right, the monomer, multimer, and complex level structures of the dARC2 capsid of *Drosophila melanogaster* [1]. The monomer and multimer are modeled with existing technologies such as AlphaFold3, OpenFold3, or Boltz-2 [2, 3, 4]. The complex is modeled with Plica-1.

After decades of incremental progress, accurate sequence-to-structure prediction was accomplished by AlphaFold2 at the monomer-level in 2020, revolutionizing structure-based drug design and discovery [7]. Soon, AlphaFold2 reached the multimer-level in 2021, opening the door to modeling and designing proteinprotein interactions and small complexes [2]. The implications for protein engineering, pharmaceuticals, and medical research in general are paradigm-altering [8].

Unfortunately, progress toward predicting structures at the protein complex level has stalled due to two fundamental issues: GPU memory constraints and the lack of experimental data on large protein complexes.

The transformer-style attention mechanisms that power AlphaFold and many other machine learning models inherently run in *O*(*n*^2^) time [2, 7, 9, 10]. Although there are approximate or restricted attention mechanisms that operate closer to linear time, performance may be sacrificed, i.e., the native sparse attention basis of DeepSeek V3.2 [11]. The quadratic scaling of these models makes it extremely difficult to build larger complexes since the memory bandwidth limit on GPUs is quickly exceeded [12]. AlphaFold3 switched to a diffusion architecture partly to compensate for the memory efficiency concern [10]. However, despite this architecture being more memory efficient, it is less compatible with parallelization operations. Altogether, this means AlphaFold3 has a size limit at ≈ 5000 amino acids, beyond which accuracy drops precipitously.

The more intractable issue is data availability. Essentially all publicly available protein structural databases in the world have already been ingested by the current AlphaFold models [2, 7, 10]. To enable progress toward large protein complexes and other important areas like antibody-receptor docking, new structural data is needed. This comes primarily from two sources: Cryo-electron microscopy (cryoEM) and X-ray diffraction (XRD), both of which are time-consuming, technically difficult, and resource-intensive procedures. These techniques can take weeks to months even in the hands of an experienced operator. Sample purification and isolation success are also heavily dependent on the target protein of interest [13]. There are likely fewer than 1000 cryoEM systems in the United States due to the significant infrastructure investment required [14]. Even if operated simultaneously at full capacity for a year, they would likely image far less than the tens of thousands of complexes required for training a new AlphaFold-like model. Experimentally generating sufficient quantities of data on large protein complexes to train new AlphaFold-like models is currently infeasible.

In the generative AI era, protein design tools such as Latent-X, RFDiffusion3, and BoltzGen are widely accessible to the community [15, 16, 17]. When provided with clear target structures, these tools, combined with the emerging rapid screening capabilities of cloud labs, will outpace traditional technologies like phage display and animal antigen immunization for antibody identification [18]. They are now bottlenecked by target discovery, i.e., selecting what proteins and where on the protein structure to target. Much of the human proteome, as well as those of viruses, animals, and plants, remains undruggable because we cannot model its structures. Therefore, there is a crucial need for modeling large protein complexes (i.e. > 60 chains and/or > 10, 000 amino acids) at computational scale in order to identify key drug targets.

With our model, which we name Plica-1, we set out to circumvent the limitations imposed by transformer architectures and experimental data availability by using physics-based heuristics. The use of such heuristics and specific internal data structures allow Plica-1 to run in *O*(*n* log *n*) time without sacrificing performance. It is also parallelizable and is not limited by hardware memory capacity. As a result, it is able to rapidly model protein complex structures far above 10,000 amino acids, with the largest demonstration at ≈40,000.

## 2 The Plica-1 model

To build a complex, Plica-1 is supplied with its symmetry type and predicted subunit structure, generated by an AlphaFold-like model. For example, for the complex in Figure 2C, Plica-1 is given its octahedral symmetry and its predicted multimer structure (in this case, a trimer). Plica-1 then assembles instances of the trimer into a cohesive complex, structuring interfaces to optimize internal parameters. The completed structure is then outputted. For a given complex, Plica-1 outputs about 5-10 solutions. The model has mechanisms which rank these solutions for physical “realness”. The predictions discussed later in this work are all the lowest root mean square deviation (RMSD) solutions, relative to experimental data, among the set of solutions outputted for each complex.

**Figure 2.**
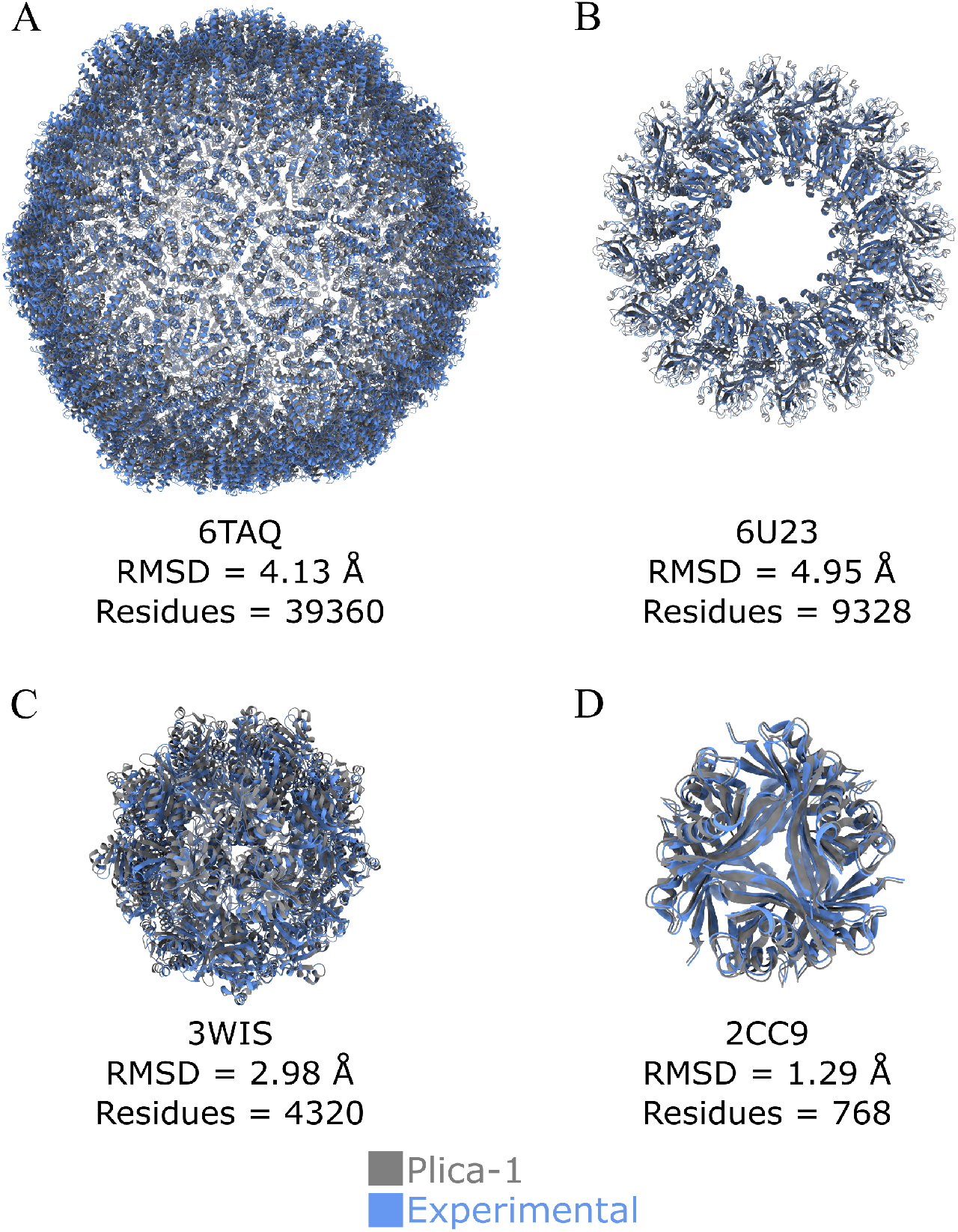
Plica-1 predicted protein complex structures overlapping with known cryoEM or X-ray crystallography data in the Protein Data Bank (PDB) [19]. **A)** The dARC2 capsid of *Drosophila melanogaster*. This structure is > 40, 000 amino acids in size, but the number of modeled residues are truncated in the cryoEM resolved structure. Iscosahedral symmetry complex. PDB code: 6TAQ [1] **B)** Human macrophage-expressed gene 1. Cyclic symmetry. PDB code: 6U23 [20] **C)** Protein cubic cage for redox transfer in *Paraburkholderia xenovorans*. Octahedral symmetry complex. PDB code: 3WIS [21] **D)** Dodecin in *Halobacterium salinarum R1*. Tetrahedral symmetry complex. PDB code: 2CC9 [22]

Existing “docking” methods use a similar approach [23, 24, 25, 26]. Such methods, whether structure-based, machine learning-based, or physics-based, serve a crucial purpose but have certain limitations. Some structure-based methods, such as RPXdock, are relatively fast and are sequence-independent [23]. This method performs best with experimentally verified or previously designed inputs. As such, publicly available literature seems to primarily focus on synthetic designs instead of naturally-occurring protein complexes, which may require more experimental data in order to model [27, 28]. A machine learning-based method, CombFold, uses AlphaFold-Multimer for combinatorial assembly of homomeric and heteromeric protein complexes [24]. This method is limited by AlphaFold-Multimer’s ability to produce pariwise subunit interactions, which, while state-of-the-art, is limited by the availability of training data, especially of protein complexes. It also has relatively long runtimes, up to ≈ 250 minutes in test cases, and, to our knowledge, has currently only been tested up to 30 protein chains and 18,000 amino acids [24]. Some prominent methods are physics-based and rely on molecular dynamics calculations. These are accurate and versatile, creating structures with accuracies on par with experimental data resolutions. The most notable of these is EvoDock, which is the nearest comparable technology to Plica-1 and is arguably the current state-of-the-art [25]. EvoDock achieves extremely high accuracies (often better than Plica-1), but requires tens to hundreds of hours of GPU time for a detailed simulation of a single, small complex. To our knowledge, it currently has not been demonstrated to build higher-order symmetric complexes such as T> 1 icosahedral capsids or cyclic complexes.

Plica-1 seeks to address the limitations discussed above with specific protein prediction models and docking methods. The use of physics heuristics instead of transformer architecture or molecular dynamics calculations makes Plica-1 extremely efficient. For these reasons, it is easily scalable to large complexes. The largest structure constructed is > 40, 000 amino acids and over 4 MegaDaltons (**Figure 1; 2A**). This is > 8-times the limits of protein prediction models like AlphaFold3, and exceeds the currently demonstrated range of molecular dynamics docking methods such as EvoDock. We eschew existing symmetry definitions used by many docking tools in favor of deriving our own symmetry library using geometric methods. This enables the modeling of naturally occurring higher-order (T > 1) icosahedral and large cyclic complexes. To our knowledge, this work represents the first fast and accurate predictions of our test cases — the human macrophage-expressed gene 1 (PDB code: 6U23) and the dARC2 capsid of *Drosophila melanogaster* (PDB code: 6TAQ) — without experimental data templates and extensive molecular dynamics calculations [1, 20].

To demonstrate the model’s capabilities, we built 36 complexes from sequence, and compared them with US-align to their experimentally measured structures in the Protein Data Bank (PDB) (**Figures 2; 3**); see appendix for a table of the complexes and their accuracies [29]. Of note, nearly all predictions had accuracies < 5Å RMSD compared to the experimental data, which makes the structures sufficiently accurate for drug design [30]. The two outliers had errors associated with the symmetry libraries which have since been identified and are currently being resolved. Using inaccurate AlphaFold-like predictions as inputs can damage the accuracy; therefore, all subunit structures used in this work had RMSD < 4Å.

About half of the predictions have accuracies within one bohr radius of the resolution of the benchmark experimental data or better (**Figure 3A**). This suggests that the model is approaching the internal resolution of the experimental methods used to generate the reference data. The accuracy of the reference data is subject to fundamental physical limits and a more accurate experimental benchmark has yet to be invented [31, 32]. Finally, we note that Plica-1’s required runtime for these predictions fol lows the expected *O*(*n* log *n*) trend (**Figure 3B**). The timed computations are for complexes featured in the EvoDock method’s publication. The same complexes which ran for tens to hundreds of hours on EvoDock were constructed by Plica-1 in minutes [25]. The Plica-1 runtimes recorded in Figure 3 were performed locally on a MacBook Air with a Apple M2 chip. Plica-1 has also been successfully parallelized to run on cloud-based virtual machines. A 96-core CPU can perform these same calculations in roughly 5-30 seconds each. This makes Plica-1 10^3^ to 10^5^ times faster than current state-of-the-art methods.

**Figure 3.**
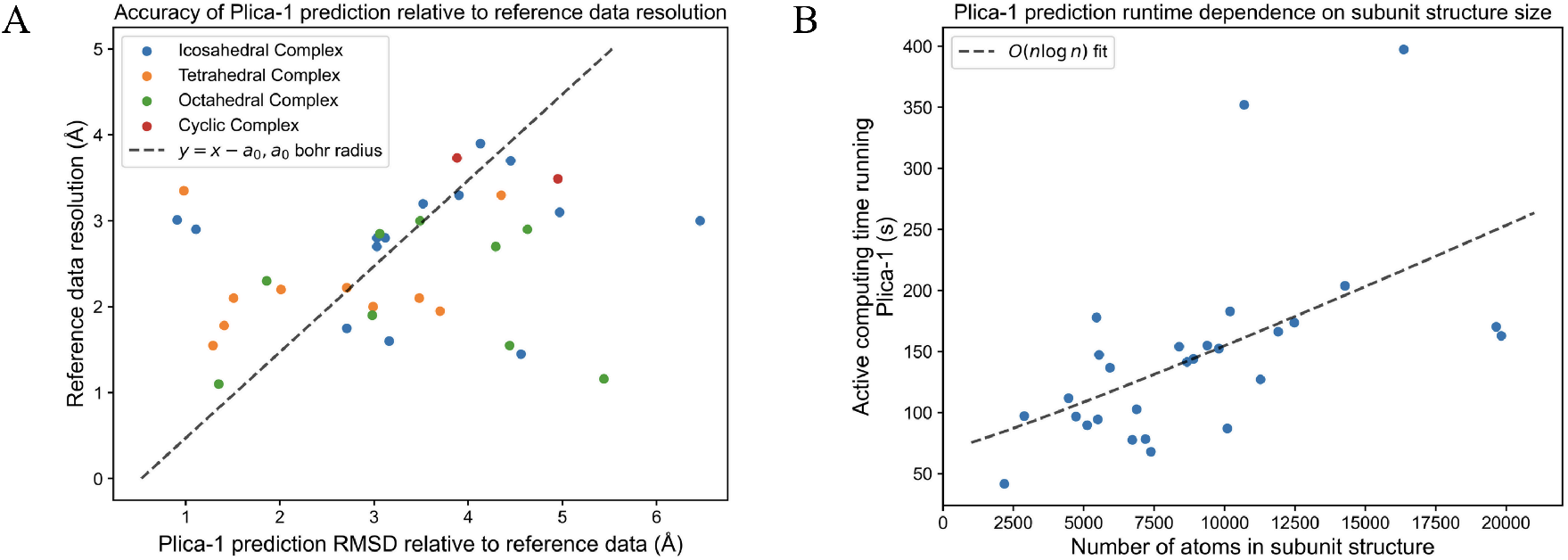
**A)** Accuracy of Plica-1 predictions with respect to cryoEM and X-ray crystallography reference data, plotted against the internal resolution of that data. **B)** Active Plica-1 prediction runtime plotted against the number of atoms in the subunit structure of the predicted complex, demonstrating an *O*(*n* log *n*) trend. After the complex is solved for, some post-processing time (< 30 seconds) is required, which includes writing the solution to a PDB file.

We note that the cyclic complexes and the T = 4 icosahedral capsid’s computing times were not included in Figure 3B. These complexes require a higher dimension parameter space and run in about 20-30 minutes on a 96-core CPU, but still run on *O*(*n* log *n*) time.

## 3 Predicting structures outside of the PDB and building a general antiviral engine

Having predicted known structures with greatly reduced computation times and with high fidelity using Plica-1, we next wanted to demonstrate the ability to predict structures not seen in the PDB, including those designed by large machine learning models. We chose Evo-Φ36, a bacteriophage designed with the Evo2 large language model, as our proof-of-concept task [33]. At the time of publication, it represented the first functional, AI-generated genome and did not have a publicly available structure in the PDB to reference. The publicly released genome and homology observations were used to annotate the Capsid F and Spike G protein of the AI designed phage. The joint pentameric structure were then predicted in AlphaFold3 (**Figure 4A**). Plica-1 then assembled the structure into a full, 3D complex with striking accuracy compared to the 2D representation of the CryoEM structure (**Figure 4A**), and ≈ 4Å RMSD relative to the wildtype phage Evo-Φ36 was engineered from. Notably, this complex is heteromeric, demonstrating that Plica-1 is not limited to homomeric complex assemblies. Prediction of the Evo-Φ36 virus points to the application of Plica-1 towards rapid antiviral design.

**Figure 4.**
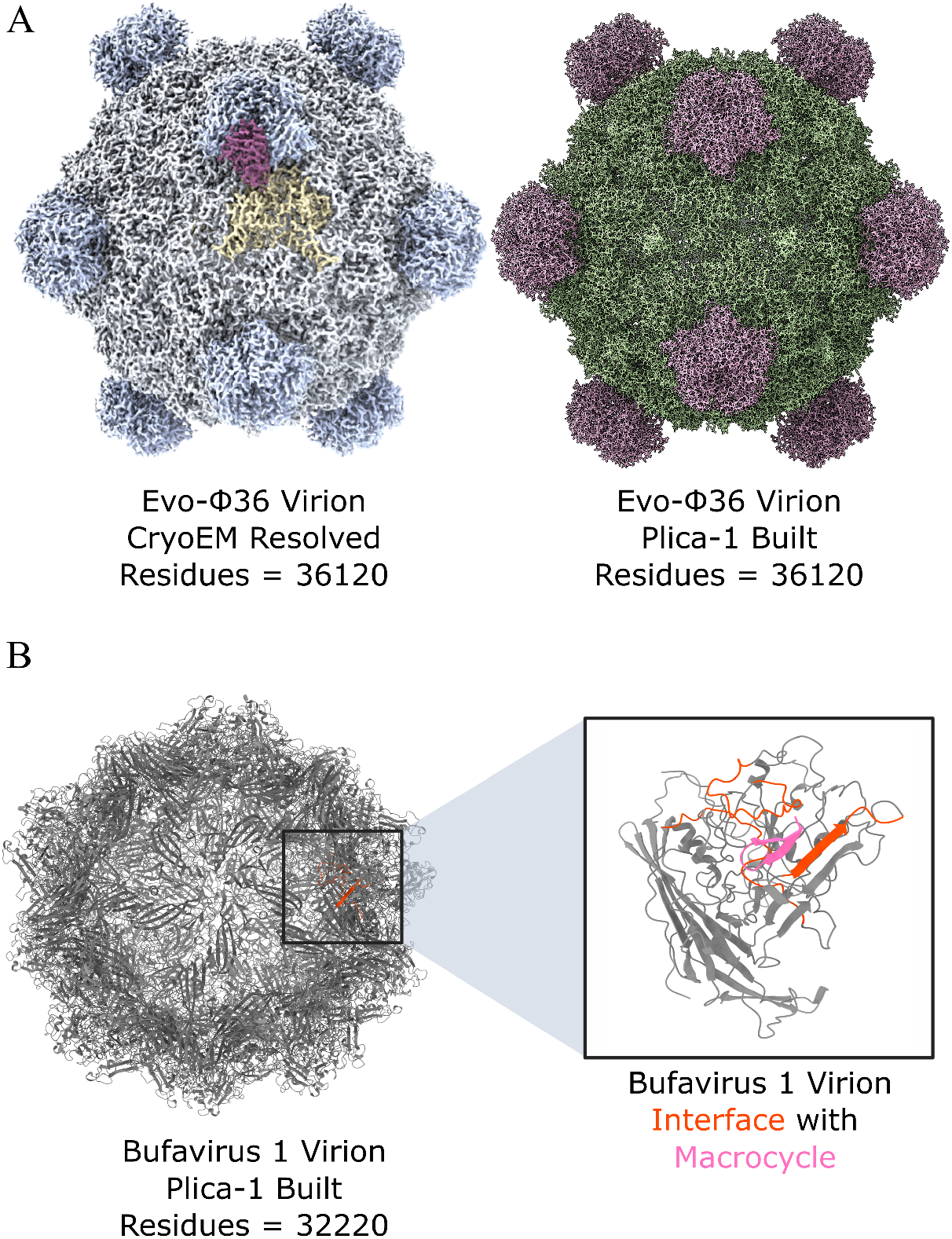
**A)** *Left*, Image of Evo-Φ36’s cryoEM structure, reproduced from Arc Institute publication [33]. *Right*, Plica-1’s prediction of Evo-Φ36 from its publicly-released genome. **B)** *Left*, Example Plica-1 prediction of Bufavirus 1 capsid structure, 6 − 7Å RMSD relative to known cryoEM datasets. *Right*, Example macrocycle inhibitor bound to Bufavirus 1 capsid, designed with Latent-X.

High resolution structural data on a virus can enable, or at least greatly facilitate, the rational design of specific antivirals targeting that virus [34, 35, 36]. New tools, such as Latent-X, RFDiffusion3, and BoltzGen, are capable of robust, semi-automated inhibitor design against protein complexes given their structure. Once a virus’s structure has been determined, antivirals can be designed against it. Unfortunately, virus structures are historically difficult to determine experimentally. In addition to all the usual challenges of cryoEM and XRD, viral pathogens work often requires a BSL-3 or even BSL-4 facility [37]. This compounded challenge often means that viral structural determination takes at least one month or, more commonly, multiple months. By the time the structure of an emerging pathogen has been experimentally determined, an outbreak may have already spread.

The most efficient method would be to predict a virus’s structure from its sequence purely computationally. This is impractical with available tools [38], but we believe Plica-1 bridges this gap. To demonstrate this capability, we used Plica-1 to predict the Bufavirus 1 capsid structure and identify sites of interaction for targeting with macrocycles to inhibit capsid formation (**Figure 4B**). Using Latent-X, we generated ≈ 300 candidates. *In vitro* validation is underway.

Combining the rapid protein complex prediction capabilities of Plica-1 with structure-based generative AI drug design models and high-throughput screening, we form the basis of a general antiviral engine. Using this workflow, a library of antiviral candidates can be designed beginning with viral sequence in under 3 hours. If sufficiently validated, this would be paradigm-altering for antiviral development. The Bufavirus 1 was chosen because it is difficult to model with Plica-1 (6 − 7Å RMSD relative to experimental data) while three high quality cryoEM structures have been reported at different pH levels [39]. This provides a reference point to help understand how AI-designed binders interact with the Plica-1-predicted structure, and how the workflow may be improved. Crucially, Bufavirus 1, to our knowledge, has no specific antiviral. A successful deployment of an antiviral in a real-world setting against Bufavirus 1 would thus be a definitive proof-of-concept.

As demonstrated painfully by the COVID-19 pandemic, essentially no country can guarantee its ability to respond quickly enough to contain an outbreak without the risk of catastrophic socioeconomic consequences. A warming and more interconnected world is at increasing risk for more global pandemics [40, 41, 42]. Beyond these naturally evolving risks, the emergence of AI-designed viruses, such as Evo-Φ36, poses an entirely new threat. Tools for designing viruses have far outpaced biosecurity infrastructure and legislation, even before the LLM era [43]. Open-access and open-source practices for leading Bio-AI tools have removed significant barriers for developing dual-use biotechnologies [44]. For both global pandemic preparedness and biodefense, there is urgent need for a universal antiviral engine.

The Plica-1-based antiviral engine is currently built for icosahedral viruses, which represent about half of known viruses [45, 46, 47]. To bridge the gap toward a universal antiviral engine, we are in the process of enabling other symmetry classes, namely helical viruses. A more difficult proposition is providing foundational models with the data they need to better model both natural and artificial virus multimers for Plica-1 to assemble and for our antiviral engine to target. A similar argument applies to the path towards universal protein structure prediction and protein complex modulation.

## 4 Toward universal protein structure prediction and protein complex modulation

Beyond viruses, we believe Plica-1’s ability to generate protein complexes *de novo* allows it to create an unrivaled synthetic dataset for training next-generation foundational models. Chai-2 demonstrated material improvement in the field of antibody-antigen engineering with the addition of synthetic data [48]. Further development of Plica-1 could provide data across protein complex classes for the next generation of models to learn. Essentially, it could create a synthetic counterpart to the PDB. This moves us towards a more complete foundational model of protein biology.

A limited, proof-of-concept version can be used to strengthen our technology for pandemic preparedness and biodefense. While AlphaFold3’s multimer predictions are state-of-the-art, they often fail for structures with insufficient homologous proteins in the PDB [49, 50]. By fine-tuning an open-source, AlphaFold-like model on a synthetic dataset of capsid complexes, we hope to improve upon capsid protein predictions.

We can then extend the model to general human health. The human proteome contains approximately 60 to 70 percent symmetric protein complexes [51]. However, the current human protein complex map, hu.MAP 2.0, has a set of ∼ 5000 complexes with more than 2 chains, and only ∼ 400 of these complexes have any reported structures [25, 52]. Complete structures exist for only a small subset of human complexes like the proteasome, ribsome, EIF2B complex, and spliceosome [52]. Furthermore, interface regions of human protein complexes show a significant enrichment of disease-causing mutations [51]. Therefore, by moving closer to modeling the entire human proteome, especially protein complex interfaces, we can unlock a substantial set of novel, druggable targets. This would be a significant step towards the eventual, field-wide goal of universal protein complex modulation: the ability to make a drug targeting any protein complex in any species by modeling its structure and designing against it. The basis of that technology would have to involve scalable prediction of large protein complex structures, and likely the synthetic data that can subsequently be generated.

In this context, it is also important to note Plica-1’s limitations and associated future directions of work. Currently, Plica-1 requires both the monomer sequence and symmetry of a complex, and models only symmetric protein complexes. While we see a clear path to important classes such as antibody-antigen complexes, there are many asymmetric complexes Plica-1 cannot model. Plica-1 has not yet been used to performed an end-to-end blind prediction: modeling a complex solely from sequence and symmetry when no experimental image of any kind has been generated. Such efforts are underway — a cryoEM image of a capsid structure is being produced, with the authors predicting its structure using Plica-1 in isolation from any experimental data. We also note that the entire Plica-1 workflow begins with high quality protein sequence. While the complex’s symmetry can often be deduced from the sequence, this is not always the case. Relying on sequence also makes the antiviral applications reactive. It cannot be deployed in advance of emerging viral threats as a preventative measure.

Meanwhile, new approaches to structural prediction that underpin Plica-1 offer opportunities for further exploration and development. With those advancements, we hope aforementioned limitations can be overcome, and new application areas discovered.

## 5 Conclusion

Plica-1 enables *de novo* prediction of large protein complex structures beyond the scale and limits of current machine learning prediction and molecular-dynamics docking methods. Operable on consumer hardware, e.g. a MacBook Air with a Apple M2 chip, it can run near-experimental resolution predictions in minutes.

Validation across symmetry classes against known structures, as well as the prediction of unknown structures, demonstrates the capability of this method in protein engineering and drug development. The ability to generate large complexes quickly, without specialized hardware, can enable applications like rapid antiviral design and generating data for training future foundational protein models.

We hope Plica-1 will serve as a valuable tool for structural biology and AI-driven drug discovery, and help move the field closer toward universal protein structure prediction and protein complex modulation.

## Acknowledgement

The authors would like to thank Rebecca Allan, Colin Maraganore, and Nelly Weiser for helpful discussions on the manuscript. We thank Seshu Ajjarapu for his mentorship and guidance. We are grateful to the facilities and operations staff at the California NanoSystems Institute for maintaining a safe and productive work space. We also acknowledge Jackson Chen for help in generating the figures in this work.

## A Plica-1 predictions data table

**Table 1:**
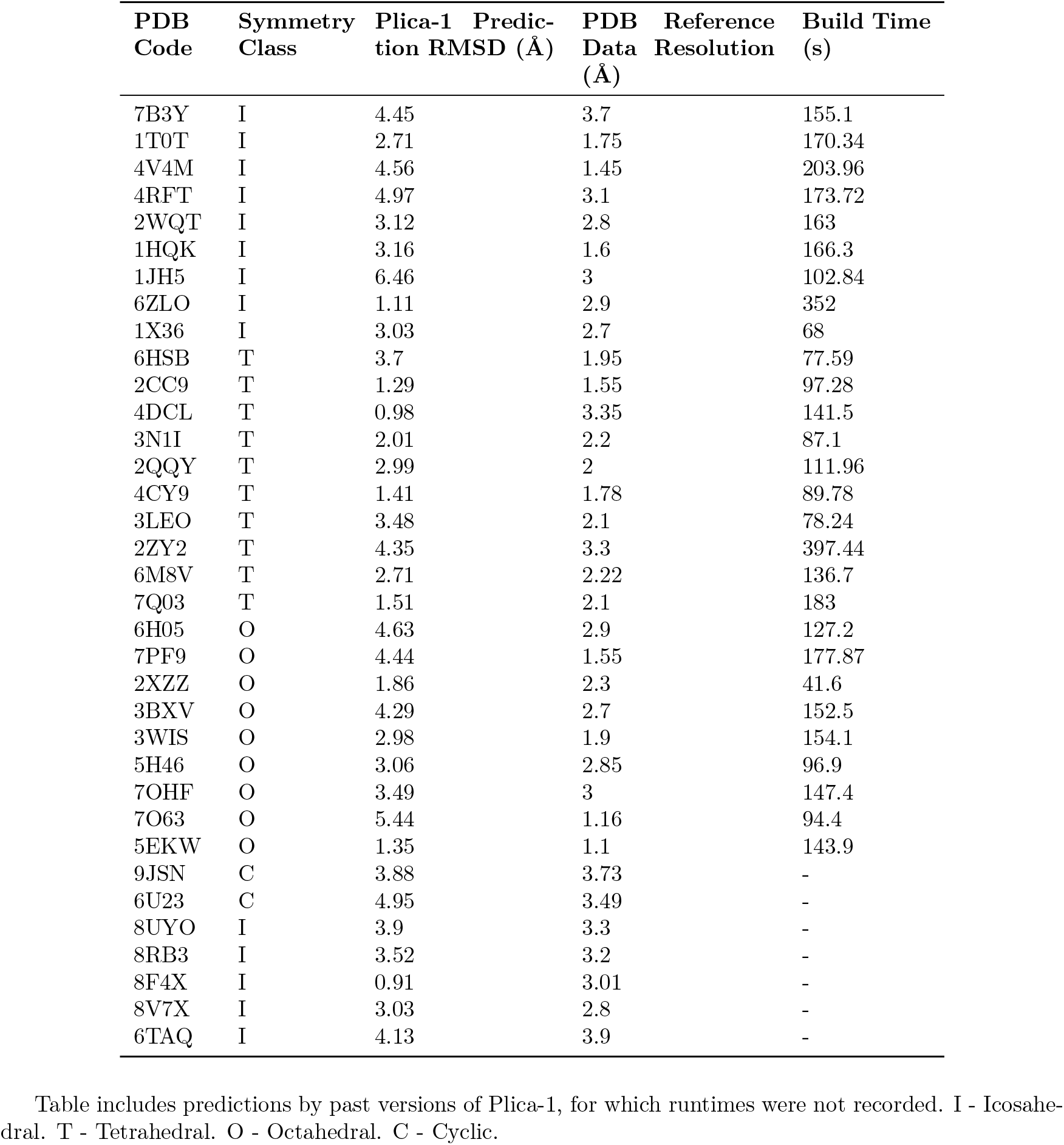
Plica-1 Protein Complex Prediction Results.

